# Retrotransposons as pathogenicity factors of the plant pathogenic fungus *Botrytis cinerea*

**DOI:** 10.1101/2021.04.13.439636

**Authors:** Antoine Porquier, Constance Tisserant, Francisco Salinas, Carla Glassl, Lucas Wange, Wolfgang Enard, Andreas Hauser, Matthias Hahn, Arne Weiberg

## Abstract

**Background:** Retrotransposons are genetic elements inducing mutations in all domains of life. Despite their detrimental effect, retrotransposons become temporarily active during epigenetic reprogramming and cellular stress response, which may accelerate host genome evolution. In fungal pathogens, a positive role has been attributed to retrotransposons when shaping genome architecture and expression of genes encoding pathogenicity factors; thus, retrotransposons are known to influence pathogenicity.

**Results:** We here uncovered a hitherto unknown role of fungal retrotransposons as being pathogenicity factors, themselves. Studying the aggressive fungal plant pathogen *Botrytis cinerea*, that is known to deliver some long-terminal repeat (LTR) deriving regulatory trans-species small RNAs (*Bc*sRNAs) into plant cells to suppress host gene expression for infection we found that naturally occurring, less aggressive *B. cinerea* strains possess considerably lower copy numbers of LTR retrotransposons and had lost retrotransposon *Bc*sRNA production. By a transgenic proof-of-concept approach, we reconstituted retrotransposon expression in a *Bc*sRNA-lacking *B. cinerea* strain, which resulted in enhanced aggressiveness in a retrotransposon and *Bc*sRNA expression-dependent manner. Moreover, retrotransposon expression in *B. cinerea* led to suppression of plant defence-related genes during infection.

**Conclusions:** We propose that retrotransposons are pathogenicity factors that manipulate host plant gene expression by encoding trans-species *Bc*sRNAs. Taken together, the novelty that retrotransposons are pathogenicity factors will have general impact on studies of host-microbe interactions and pathology.

## Background

Retrotransposons are genetic elements inducing mutations in all domains of life [1]. In eukaryotes, distinct classes of suppressive, cis-regulatory sRNAs, such as PIWI-associated piRNAs in Drosophila and nematodes [2] and heterochromatic small interfering RNAs in plants, are produced for retrotransposon control [3, 4]. Despite their detrimental effect, retrotransposons become temporarily active during epigenetic reprogramming [5] and cellular stress response [6, 7], which may accelerate host genome evolution [8–11]. In fungal pathogens, a positive role has been attributed to retrotransposons to shaping genome architecture and expression of genes encoding pathogenicity factors [12, 13]; thus, retrotransposons can influence pathogenicity.

*B. cinerea* can infect > 1400 plant species, including important crops such as tomato [14, 15]. For infection, *B. cinerea* small RNAs (*Bc*sRNAs) translocate into plants and hijack the plant Argonaute (AGO)1/RNA-induced silencing complex (RISC) to suppress host immunity genes. Cross-kingdom sRNA effectors have also been reported in other fungal, oomycete, bacterial, and parasitic plant species [16–19], making cross-kingdom RNA interference (ckRNAi) a remarkably common phenomenon in plant-pathogen interactions [20]. The vast majority of *Bc*sRNA effectors derive from long-terminal repeat (LTR) retrotransposons [21, 22], raising the hypothesis that retrotransposons could play a role as fungal pathogenicity factors.

### Botrytis cinerea strains carrying LTR retrotransposons are more aggressive

Previous transposon annotation in a published *B. cinerea* genome release (strain B05.10) revealed 83 full-length LTR retrotransposon copies separated into nine different consensus classes (Fig.S1) either belonging to the Gypsy or the Copia superfamily [23] (Tab.S1). Using this *B. cinerea* LTR retrotransposon collection, we defined six subfamilies according to phylogenetic analysis, which we named *Bc*Gypsy1-*Bc*Gypsy4 and *Bc*Copia1-*Bc*Copia2 (Fig.1A, Tab.S1, S2). We analysed previously published sRNA-seq data (raw data are available at NCBI GEO: GSE45323, GSE45321) regarding LTR retrotransposon *Bc*sRNA production in *B. cinerea* axenic culture grown on agar plates or during infection of the host plants *Solanum lycopersicum* (tomato) and *Arabidopsis thaliana* [22]. LTR retrotransposon *Bc*sRNA production in axenic culture indicated that only *Bc*Gypsy1, *Bc*Gypsy3 and *Bc*Gypsy4 produced significant amounts of *Bc*sRNAs (Fig.S2A). During plant infection, more than 80% of LTR retrotransposon *Bc*sRNAs mapped to *Bc*Gypsy3. These *Bc*sRNAs displayed induced accumulation at early infection time point (Fig.1B, Fig.S2B) in contrast to *Bc*sRNAs derived from other genomic loci (Fig.S2C) suggesting a role in pathogenicity.

**Figure 1:**
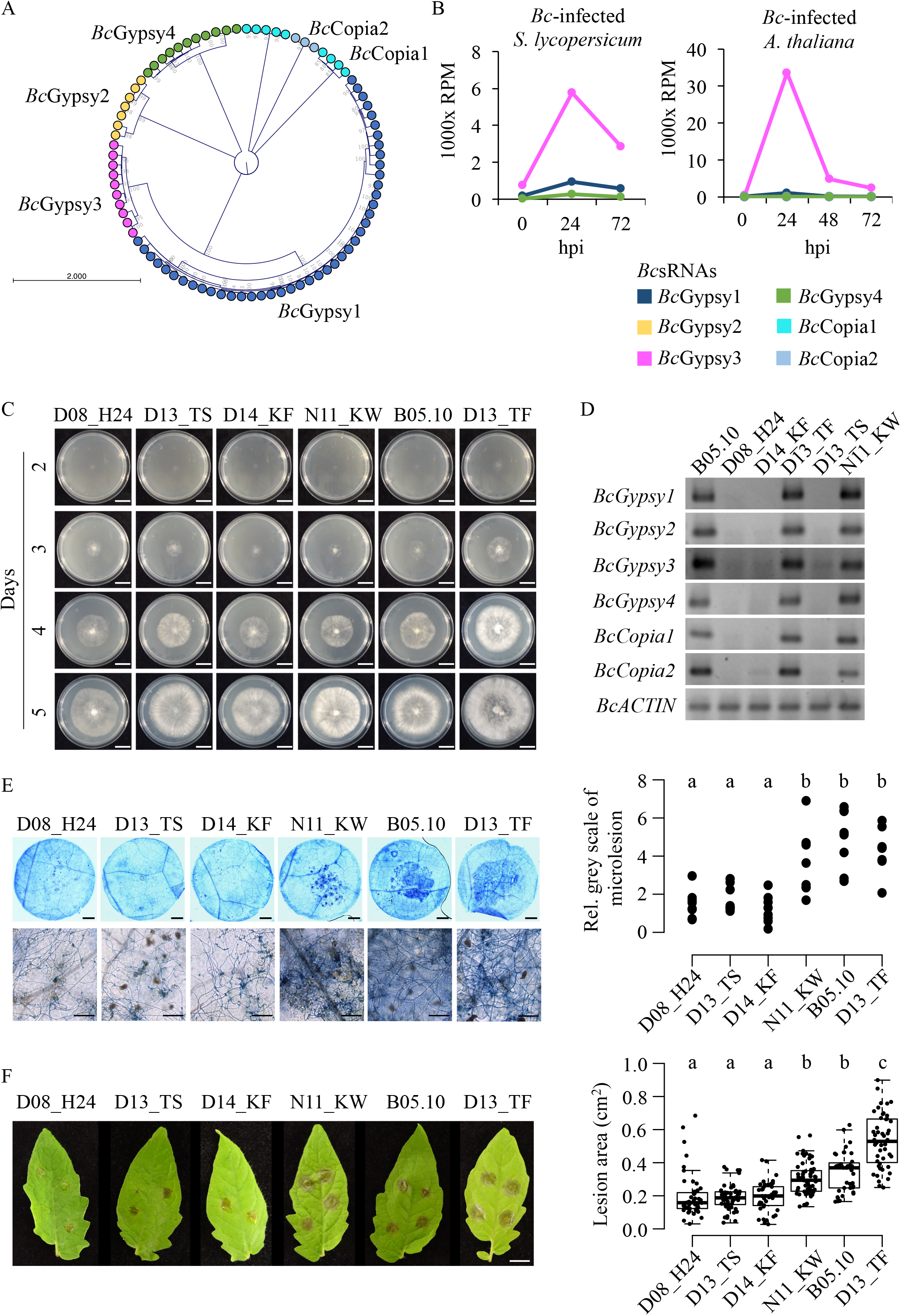
*B. cinerea* strains carrying LTR retrotransposons are more aggressive. A) LTR retrotransposon phylogenetic relationship of six subfamilies, *Bc*Gypsy1-4, *Bc*Copia1-2, identified in the strain B05.10. B) LTR retrotransposon *Bc*sRNA abundance of the six subfamilies (*Bc*Gypsy1-4, *Bc*Copia1-2) at different time points of *S. lycopersicum* or *A. thaliana* host infection (raw data are available at NCBI GEO: GSE45323, GSE45321). C) Growth phenotype of the six *B. cinerea* strains. Scale bars indicate 20 mm. D) Genotyping PCR of the six LTR retrotransposon subfamilies in the six *B. cinerea* strains. E) Pathogenicity assay of the six *B. cinerea* strains on tomato leaves quantifying microlesion area by Trypan Blue staining at 24 hpi. Microlesions in Trypan Blue images were quantified in eight leaf discs per strain as relative grey scale (rel. counts) in relation to the total leaf disc. Scale bars in leaf disc image indicate 1 mm, and in higher magnification images 100 μm. F) Pathogenicity assay of the six *B. cinerea* strains on tomato leaves indicating lesion area of > 20 infection sites at 48 hpi. Scale bar indicates 1 cm. For F), infection experiments were repeated at least three times with similar results. In E) and F), significant difference is indicated by letters, and was tested by one-way ANOVA using Tukey HSD test with p< 0.05.

To investigate the contribution of LTR retrotransposons to the pathogenicity of *B. cinerea*, we analysed six *B. cinerea* strains collected from different host plants and geographical origins (Tab.S3) at genomic, sRNA transcriptomic, and disease phenotypic levels. Strains grown on nutrient-rich agar showed no obvious phenotypic differences; however, the strain D13_TF grew faster (Fig.1C, Fig.S3A) and did not form sclerotia (Fig.S3B), a dormant tissue that enables *B. cinerea* to survive non-favourable conditions. We designed primers to genotype the six *B. cinerea* strains for the genomic presence of the six LTR retrotransposon subfamilies by PCR. The strains D08_H24, D14_KF and D13_TS were negatively tested for all LTR retrotransposon subfamilies. In contrast, B05.10, N11_KW and D13_TF were tested positive for all subfamilies (Fig.1D). We performed pathogen assays with the six *B. cinerea* strains on tomato leaves to assess aggressiveness. We defined strain aggressiveness as a quantitative level of its ability to induce disease symptoms in the infected host. Tomato was chosen as a suitable host plant, as it was found to be susceptible to LTR retrotransposon *Bc*sRNAs triggering ckRNAi [22]. The three strains negatively tested for LTR retrotransposons were less aggressive compared to the three LTR retrotransposon positively tested strains, considering induction of primary lesion formation at 24 h post inoculation (hpi) (Fig.1E) and extended lesion area on infected leaves at 48 hpi (Fig.1F). Based on these results, we found a positive relationship between LTR retrotransposons and *B. cinerea* aggressiveness.

### Aggressive B. cinerea strains produce massive retrotransposon small RNAs

To gain further insights into the relationship between *B. cinerea* aggressiveness and LTR retrotransposon *Bc*sRNAs, we performed a comparative sRNA-seq analysis using *B. cinerea* cultures. Remarkably, genome-wide *Bc*sRNA mapping revealed that only the three most aggressive strains produced massive amounts of transposon-derived *Bc*sRNAs (Fig.2A). After filtering out rRNA reads, we further quantified relative read numbers of *Bc*sRNAs aligning to annotated transfer (t)RNAs, small nuclear/nucleolar (sn/sno)RNAs, protein coding genes (mRNAs) or transposons in all six *B. cinerea* strains. Consistently, the three most aggressive strains produced high amounts of transposon-derived *Bc*sRNAs between 2 - 16%, compared to 0.03 - 0.12% by the three less aggressive strains (Fig.2B, Tab.S4). In this analysis, transposon-derived *Bc*sRNAs mostly mapped to LTR retrotransposons (>99.4% in the most aggressive strains), leaving *Bc*sRNAs that derived from DNA transposons (Tab.S2) of minor proportion (Tab.S4). Size profiles of total *Bc*sRNAs from the six strains were rather diverse (Fig.2C), but LTR retrotransposon *Bc*sRNAs showed a clear size preference of 21-22 nucleotides (nt) and preferentially 5` terminal Uracil (U) (Fig.2D). In plants, AGO1 commonly associates with endogenous 21-22 nt and 5`U sRNAs to form an AGO1/RISC [24], which likely explains why *B. cinerea* LTR retrotransposon *Bc*sRNAs bind to the plant AGO1 and induce ckRNAi [21, 22]. In this regard, the majority of LTR retrotransposon *Bc*sRNA effectors (29 out of 48) mapped to *Bc*Gypsy1 and *Bc*Gypsy3 subfamilies. In accordance, the less aggressive strains D08_H24, D14_KF and D13_TS showed nearly no expressed *Bc*sRNAs aligning to *Bc*Gypsy1 and *Bc*Gypsy3, whereas many were found in the aggressive B05.10, N11_KW and D13_TF strains (Fig.S4). The strain D13_TF produced less LTR retrotransposon *Bc*sRNA compared to B05.10 and N11_KW (Fig.2A-B, Fig. S4), however it induced in average the biggest lesions (Fig.1F). The relative large lesion formation could be explained with its fast growth (Fig.1C, Fig.S3A). With these results, we found evidence that the less aggressive *B. cinerea* strains lacked production of LTR retrotransposon *Bc*sRNAs, which further supported that LTR retrotransposon *Bc*sRNAs are important for pathogenicity.

**Figure 2:**
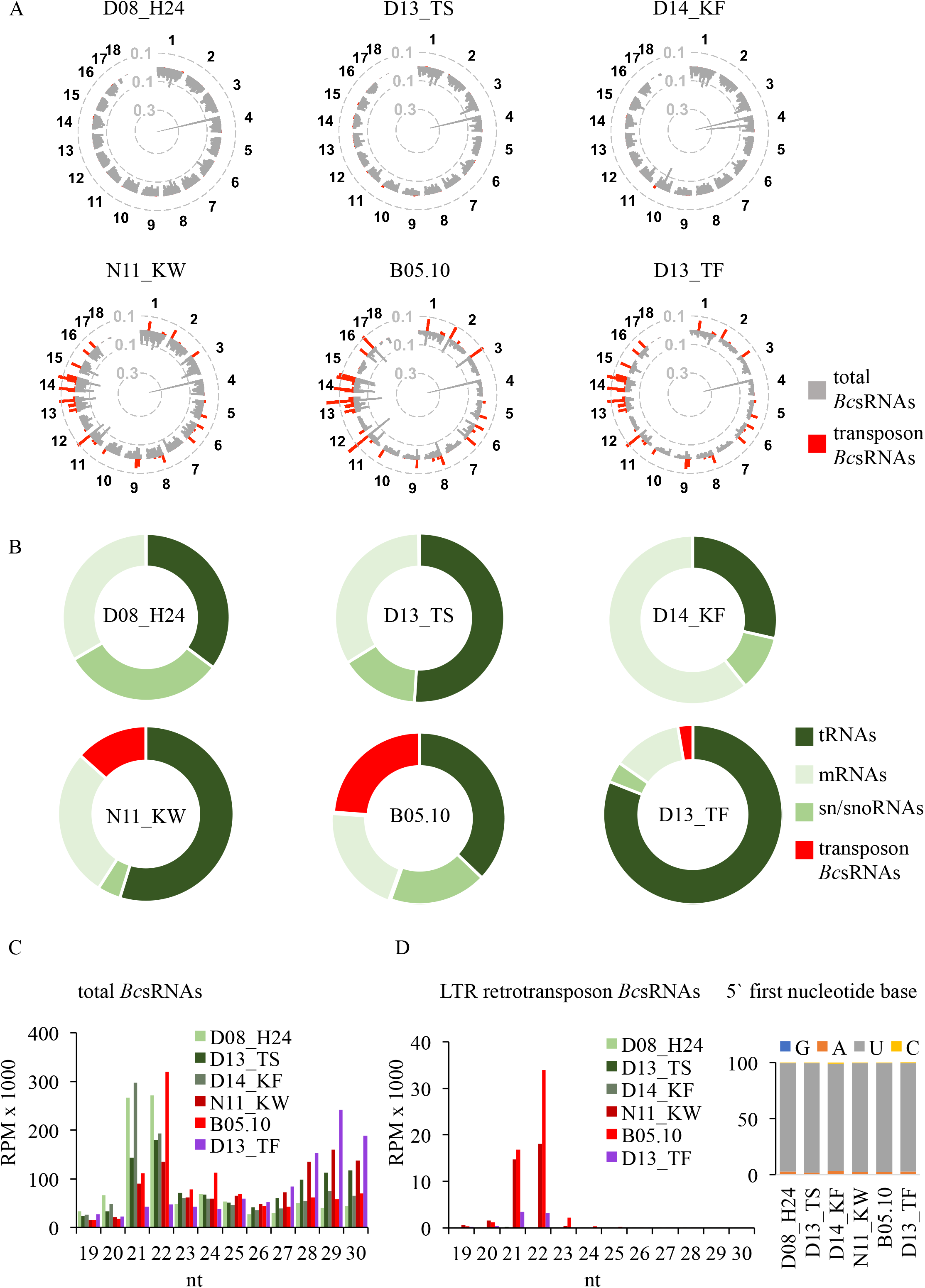
Comparative sRNA-seq analysis of the six *B. cinerea* strains. A) Genome-wide *Bc*sRNA maps of the six *B. cinerea* strains, coverage represented as log(RPM). B) Relative composition of *Bc*sRNAs mapping to distinct genomic loci. C) Size profiles of *Bc*sRNAs in the six *B. cinerea* strains. D) Sizes profiles and 5` first nucleotide distribution of LTR retrotransposon *Bc*sRNAs in the six *B. cinerea* strains.

### Less aggressive B. cinerea strains carry only silenced LTR retrotransposons relics

To clarify if the reason for the absence of LTR retrotransposon *Bc*sRNAs in less aggressive *B. cinerea* strains was due to the lack of LTR retrotransposons, as indicated by genotyping PCR (Fig.1D), we re-sequenced the genomes of the two less aggressive strains D08_H24 and D14_FK using hybrid Nanopore long-read in combination with Illumina short-read sequencing. Scaffold assembly resulted in 49 contigs for D08_H24 and 28 contigs for D14_KF. Pairwise genome alignment and synteny analysis of D08_H24 and D14_KF with the published B05.10 genome [25] revealed coverage of all 18 chromosomes (Fig.3A). The nearly gapless full genome assemblies of D08_H24 and D14_KF allowed us to perform transposon annotation and analysis. Using the REPET pipeline [26] on the D08_H24 and D14_KF genomes, we did find no Gypsy LTR retrotransposon and only one Copia LTR retrotransposon with 14 and 10 full-length copies, respectively (Fig.3B, Tab.S5). To complement our REPET analysis, we performed Blastn search in the published B05.10, and the D08_H24 and D14_KF re-sequenced genomes, using REPET-annotated full-length LTR retrotransposons of B05.10 as queries and allowing a minimum alignment length of 400 nucleotides (nt). By this step, we identified further Gypsy and Copia elements in B05.10, D08_H24, and D14_KF (Fig.3B), with fewer and more truncated ones in D08_H24 and D14_KF compared to in B05.10 (Fig.3C, Fig.S5A-B). Using all retrotransposon copies annotated in B05.10 and the re-sequenced D08_H24 and D14_KF genomes either identified by REPET or Blastn for sRNA-seq read mapping, we confirmed that D08_H24 and D14_KF produced marginal amounts of LTR retrotransposon *Bc*sRNAs compared to B05.10 (Fig.3B, Tab.S6). A striking difference regarding LTR retrotransposons between D08_H24, D14_KF, and B05.10 was the GC content. While D08_H24 and D14_KF carried exclusively full-length retrotransposons with low GC content (<30 %), B05.10 displayed two distinct fractions of low GC (<30 %) or high GC (>40 %) contents (Fig.3D, Tab.S2, S5). We obtained similar results for truncated retrotransposon copies identified by Blastn, although few truncated copies in D08_H24 and D14_KF showed GC content >40% (Fig.S6). Low GC content let us presume that LTR retrotransposons were mutated by the Repeat-induced point mutation (RIP), which is a fungal genome surveillance mechanism mutating C to T bases within duplicated sequences [27]. Previous RIP analysis in *B. cinerea* had indicated bias for CA to TA and CT to TT dinucleotides [28]. In order to estimate the occurrence of RIP on REPET-detected full-length retrotransposons, we calculated the TA/AT index. We found diverse TA/AT ratios in B05.10, but only high (>0.89) TA/AT index in D08_H24 and D14_KF (Fig.3E, Tab.S7), a threshold considered to indicate RIP [29]. Similarly, comparative analysis of both full-length and truncated retrotransposon copies in B05.10, D08_H24 and D14_KF using the RIPCAL program [29] revealed RIP in all copies of D08_H24 and D14_KF, but some copies in B05.10 with low RIP, as exemplified for the *Bc*Gypsy3 subfamily (Fig.S7). All full-length LTR retrotransposons found in D14_KF or D08_H24 displayed high RIP index and low GC contents (<30%). This finding possibly explains why LTR retrotransposon genotyping PCR was negative for D08_H24 and D14_KF (Fig.1D), as primer sequences did not match the mutated LTR retrotransposon sequences. In fungi, RIP often leads to transcriptional silencing [30], and we anticipated that loss of LTR retrotransposon *Bc*sRNAs could be a result of low LTR retrotransposon expression. Indeed, expression values of *Bc*sRNAs mapping to *Bc*Gypsy1, *Bc*Gypsy3 and *Bc*Gypsy4 correlated positively with the LTR retrotransposon GC content and negatively with their TA/AT index (Fig.3F, Tab.S8) in B05.10. GC-rich copies of *Bc*Gypsy2, *Bc*Copia1 and *Bc*Copia2 showed low *Bc*sRNA mapping accumulation in general; still, more reads mapped to GC-rich copies within the corresponding LTR retrotransposon subfamily (Tab.S9). Our genome-wide comparative analysis of LTR retrotransposons revealed that less aggressive *B. cinerea* strains possessed fewer copy numbers with low GC content, which did not produce LTR retrotransposon *Bc*sRNAs.

**Figure 3:**
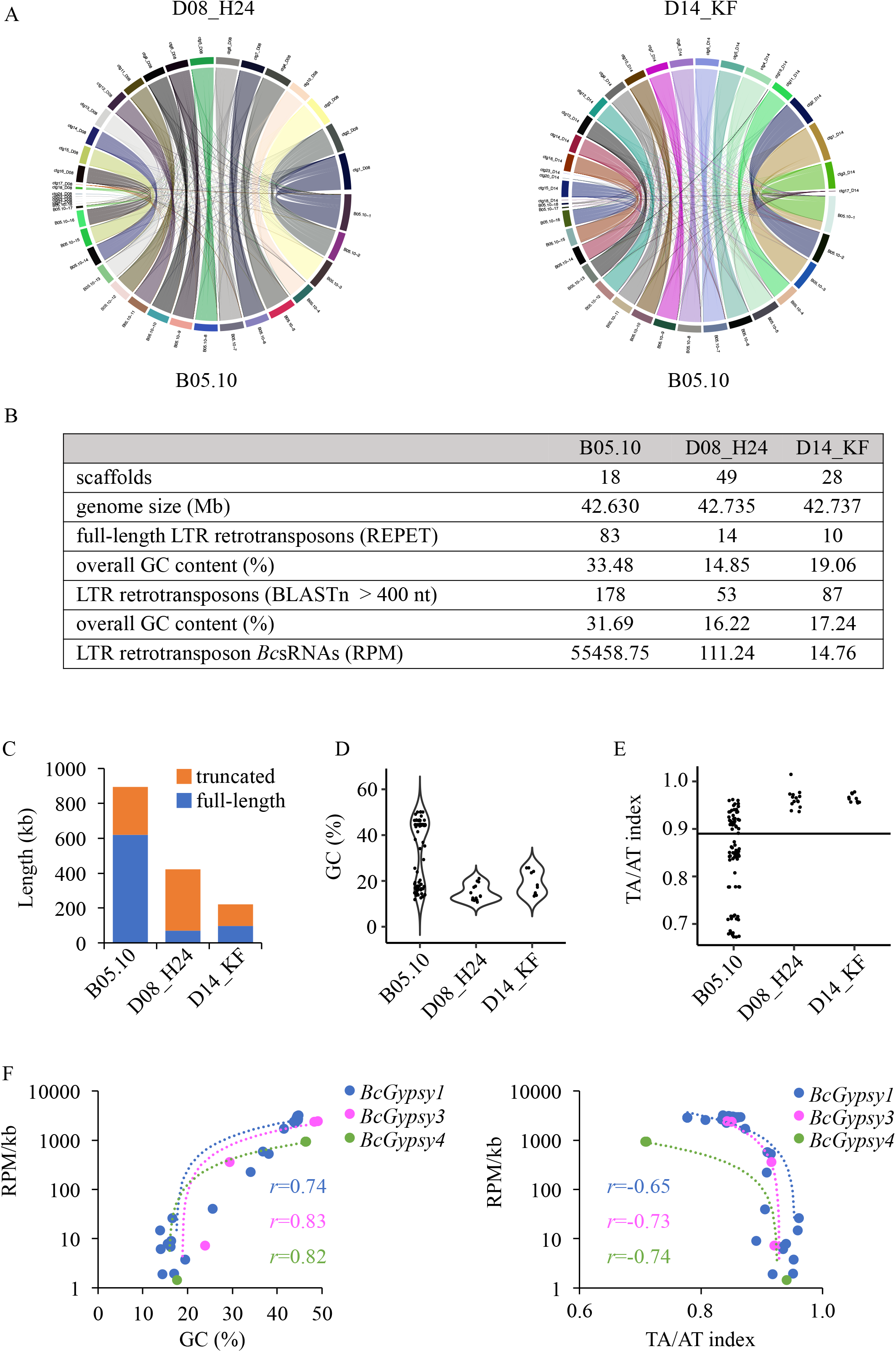
Comparative genome analysis of LTR retrotransposons. A) Whole genome chromosomal alignment analysis. B-E) Comparative analysis of LTR retrotransposons identified in the strains B05.10, D08_H24, and D14_KF, and differences in the level of copy numbers, truncation, GC content (%), and the TA/AT dinucleotide ratio with threshold line shown at 0.89. F) Correlation analysis between LTR retrotransposon *Bc*sRNA read abundance and GC content (%) or TA/AT dinucleotide ratio of the strain B05.10. *r* gives the Pearson correlation coefficient.

### The B. cinerea retrotransposon BcGypsy3 promotes host plant infection

Having observed a positive relationship between full-length, GC-rich LTR retrotransposons, LTR retrotransposon *Bc*sRNAs, and pathogenicity, we sought to validate the concept of retrotransposon as pathogenicity factor by a transgenic approach. We cloned a GC-rich *Bc*Gypsy3 from the aggressive B05.10 strain to transform the less aggressive D08_H24, anticipating that transformed D08_H24 would produce *Bc*Gypsy3 *Bc*sRNAs and gain pathogenicity. We chose a GC-rich *Bc*Gypsy3 as a major source of *Bc*sRNAs, and chose D08_H24, because this strain did not possess any full-length *Bc*Gypsy3 (Fig.S5B, Fig.S8) and did not produce significant amounts of LTR retrotransposon *Bc*sRNAs (Fig.2, Tab.S6). A 4.98 kilobase *BcGypsy3* was cloned into a fungal expression vector. We excluded flanking LTR DNA from cloning, because these typically contain binding sites of tRNA primers that initiate reverse transcription and transposon transposition [31, 32], and because *Bc*sRNAs did not derive from the LTR regions (Fig.S2B). We generated two *Bc*Gypsy3 cassettes driven by constitutive promoters to either produce single strand (ss)*Bc*Gypsy3 sense RNAs or sense and antisense RNAs to form double strand (ds)*Bc*Gypsy3 RNAs to enhance *Bc*sRNA production. Empty vector (EV) was used as transformation control (Fig.S9A). Transgenes were inserted into the *Nitrate reductase D* (*BcniaD)* locus by homologous recombination, which was previously established for targeted transgene insertion in *B. cinerea* without affecting pathogenicity [33]. Genomic insertion and *BcGypsy3* mRNA expression were validated by PCR (Fig.S9B) and quantitative reverse transcription (qRT)-PCR (Fig.4A), respectively. We performed pathogen assays with *Bc*Gypsy3 transformants that grew comparably on nutrient-rich agar (Fig.S9C) to test for increased aggressiveness. Four independent ds*BcGypsy3* transformants induced significantly larger lesions than an EV transformant (Fig.4B). Replicating this experiment with four ss*Bc*Gypsy3 transformants gave similar results, albeit gained pathogenicity was not as strong as in the ds*BcGypsy3* transformants (Fig.4B). These results were in line with a higher *BcGypsy3* RNA expression found in the ds*Bc*Gypsy3 transformants compared to the ss*Bc*Gypsy3 transformants (Fig.4A). We selected one ss*Bc*Gypsy3 (transformant #51), one ds*Bc*Gypsy3 (transformant #56) and the EV strain (transformant #18) for sRNA-seq analysis to relate *Bc*Gypsy3 *Bc*sRNA production to pathogenicity. *Bc*Gypsy3 *Bc*sRNA accumulation was moderately higher in the ds*Bc*Gypsy3 compared to the ss*Bc*Gypsy3 (Fig.4D), which was in line with the higher *BcGypsy3* RNA expression and aggressiveness of ds*Bc*Gypsy3 transformants. However, overall *Bc*Gypsy3 *Bc*sRNA accumulation in the transgenic D08_H24 was not as high as in B05.10, although the ds*Bc*Gypsy3 transformants displayed similar *BcGypsy3* RNA expression (Fig.4A). We therefore suspected that insertion of the *BcGypsy3* transgene into the *BcniaD* locus might have limited *Bc*sRNA production. To test this possibility, we repeated transformation without *BcniaD* homologous recombination flanking DNA for *BcGypsy3* random genome integration. We isolated two independent, random-inserted *Bc*Gypsy3 (rand*Bc*Gypsy3) transformants #1, #3 with confirmed *BcGypsy3* RNA expression (Fig.4A) and exhibiting comparable growth to the other isolated transformants (Fig.S9C). The rand*Bc*Gypsy3 #1 was significantly more aggressive compared to D08_H24 wt and displayed highest *Bc*Gypsy3 RNA expression (Fig.4A, C). Sequencing sRNAs of the rand*Bc*Gypsy3#1 revealed that production of *Bc*Gypsy3 *Bc*sRNAs was at approximately 10 times higher than in the transformants with *BcGypsy3* inserted into the *BcniaD* locus (Fig.4D, Fig.S10), which unexpectedly correlated with the highest *Bc*Gypsy3 mRNA expression in the rand*Bc*Gypsy3 #1 (Fig.4A). However, *Bc*Gypsy3 *Bc*sRNA production in the rand*Bc*Gypsy3 #1 was around 35 times lower than in B05.10 (Fig.4D). Taken together, transgene-induced *Bc*Gypsy3 *Bc*sRNA expression led to enhanced aggressiveness, which strongly supported that this retrotransposon is a pathogenicity factor in *B. cinerea*.

**Figure 4:**
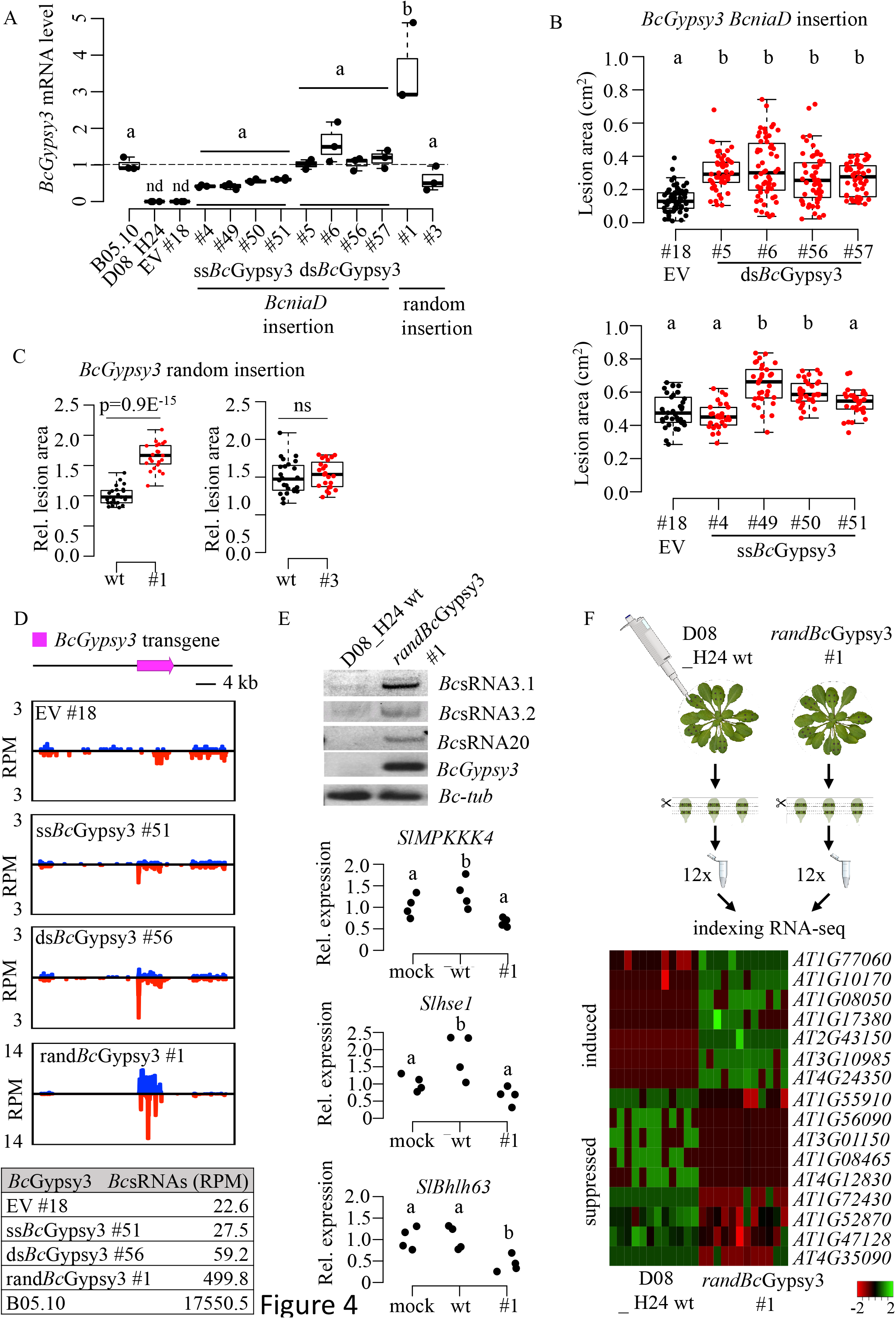
*Bc*Gypsy3 promotes host plant infection. A) *BcGypsy3* RNA expression in single strand (ss)*Bc*Gypsy3 (#4, #49, #50, #51), double strand (ds)*Bc*Gypsy3 (#5, #6, #56, #57) or empty vector (EV) (#18) transformants with the *BcGypsy3* transgene inserted in the *BcniaD* locus of D08_H24. Further, randomly inserted ss*Bc*Gypsy3 (rand*Bc*Gypsy3) tranformants #1, #3 into the D08_H24 genome, as well as the wild type strains D08_H24 and B05.10 are shown. Data points represent biological replicates. B) Pathogenicity assay with dropped spore suspension of *Bc*Gypsy3 transformants on tomato leaves quantifying lesion area of > 20 infection sites at 48 hpi. Significant difference is indicated by letters, and was tested by one-way ANOVA using Tukey HSD test with p< 0.05. C) Pathogenicity assay with agar plugs of rand*BcGypsy3* transformants #1 and #3 on tomato leaves quantifying lesion area of minimum 8 infection sites at 48 hpi. Infection series were repeated three times with similar results. Significant difference was tested by two-sided Student`s t-test. For B) and C), similar results were obtained in two independent infection experiments. D) *Bc*sRNA maps from sRNA-seq analysis at the *BcGypsy3* transgene locus representing ss*Bc*Gypsy3 #51, ds*Bc*Gypsy3 #56, rand*Bc*Gypsy3 #1 or EV #18 transformants with blue bars indicate sense and red bars antisense reads. Table shows normalized read counts (RPM) of *Bc*Gypsy3 *Bc*sRNAs. E) On top, Stem-loop RT-PCR showed *Bc*sRNA3.1, *Bc*sRNA3.2 and BcsRNA20 expression in the rand*Bc*Gypsy3 #1. *BcGypsy3* mRNA expression and *BcTub* were used as controls. On the bottom, qRT-PCR of *S. lycopersium* target mRNA expression after water treatment (mock) or after infection with D08_H24 wt or rand*Bc*Gypsy3 #1. Each data point represents a biological replicate. F) On the top, scheme of RNA-seq experiment infecting *A. thaliana* either with D08_H24 wt or rand*BcGypsy3* #1 comprising 12 biological replicates for each treatment. On the bottom, heat map showing differentially expressed *A. thaliana* genes comparing infection with D08_H24 wt or rand*Bc*Gypsy3 #1. Scale bars in B) and C) represents 1 cm.

We hypothesized that the *Bc*Gypsy3 transformants exhibited enhanced aggressiveness due to production of *BcGypsy3 Bc*sRNAs suppressing plant target genes. To test this hypothesis, we chose three known *Bc*Gypsy3 *Bc*sRNA effectors, *Bc*sRNA3.1, *Bc*sRNA3.2 and *Bc*sRNA20, and their target genes in tomato (Fig.S11A) [22]. We confirmed expression of *Bc*sRNA3.1, *Bc*sRNA3.2 and *Bc*sRNA20 in the rand*Bc*Gypsy3 #1 by stem-loop RT-PCR (Fig.4E). We then infected plants either with rand*Bc*Gypsy3 #1 or D08_H24 wt in comparison to mock-treated plants and measured mRNA levels of the tomato *Mitogen-activated protein kinase kinase kinase 4* (*SlMPKKK4*), *Class E vacuolar protein-sorting machinery protein hse1* (*Slhse1, Solyc09g014790*) and the *Basic helix-loop-helix (Bhlh)63 transcription factor* (*Solyc03g120530*). The tomato target genes showed down-regulation upon infection with rand*Bc*Gypsy3 #1 compared to D08_H24 wt (Fig.4E). To repeat this experiment with a different plant species, we infected *A. thaliana* with D08_H24 wt or rand*Bc*Gypsy3 #1 to quantify gene expression of the known *Bc*sRNA3.1 and *Bc*sRNA3.2 target genes *AtMPK1*, *AtMPK2* and a *Cell wall-associated kinase* (*AtWAK*) (Fig.S11B) [22]. *AtMPK1* showed significant down-regulation, and *AtMPK2* in tendency, upon infection with rand*Bc*Gypsy3 #1 compared to D08_H24 wt (Fig.S11C). The *A. thaliana* infection-responsive genes *AtPlant Defensin(PDF)1.2 and AtPathogensis-related protein (PR)1,* which were all not predicted targets of *Bc*sRNAs, did not show downregulation in plants infected with rand*Bc*Gypsy3 #1 compared to D08_H24 wt (Fig.S11C) indicating that plant gene downregulation was not a general effect of rand*Bc*Gypsy3 infection. The downregulation of target genes supported the role of *Bc*Gypsy3 *Bc*sRNAs plant gene manipulation.

To further explore the impact of the *BcGypsy3* transgene on the plant mRNA transcriptome during infection, we conducted an RNA-seq experiment comparing *A. thaliana* plants infected with rand*Bc*Gypsy3 #1 or D08_H24 wt (Fig.4F). Differential gene expression analysis of *A. thaliana* mRNAs indicated only moderate effects on a small subset of *A. thaliana* genes. In total, we identified 7 up- and 8 down-regulated candidate genes when plants were infected with the rand*Bc*Gypsy3 #1 by differential expression analysis applying a False discovery rate (FDR) cut-off at 0.05 (Fig.4F, Tab.S10). In consistence, similar pattern of up- or downregulation was evident for the candidate genes according to the Bio-Analytic Resource for Plant Biology (BAR) database [34] (Fig.S12). Down-regulated genes included the *Small auxin upregulated RNA 78* (*AT1G72430*) and the *Catalase (CAT)2* (*AT4G35090*), both related to auxin signalling. Repression of auxin response was previously shown to increase *A. thaliana* susceptibility to *B. cinerea* [35]. *CAT2* as well as the downregulated cysteine protease *Response to dehydration 21* (*AT1G47128*) are *A. thaliana* resistance factors against *B. cinerea* [36, 37]. Up-regulated genes related to stress response included the transcriptional repressor *NF-X-LIKE 1* (*At1G10170*) and the transcriptional repressor *Jasmonate-Zim-Domain* (*JAZ)5* (*At1G17380*), both being negative regulators of *A. thaliana* defence response against fungal phytotoxins [38] and *B. cinerea* [39], respectively. Thus, *Bc*Gypsy3 contributed to manipulating plant gene expression, possibly via *Bc*Gypsy3 *Bc*sRNA-induced ckRNAi. In this sense, we predicted 5 out of the 7 rand*Bc*Gypsy3 #1 down-regulated *A. thaliana* genes as targets of *Bc*Gypsy*3 Bc*sRNAs by psRNATarget [40], but also found predicted *Bc*sRNA targets among the up-regulated *A. thaliana* genes suggesting both direct and indirect effects on plant gene regulation (Tab.S11).

## Discussion

Our data provide lines of evidence that retrotransposons are pathogenicity factors in *B. cinerea*. Previous observation suggested that *B. cinerea* strains containing transposons colonize more frequently host plants and might induce stronger disease severity compared to transposon-free strains [41, 42]. In this regard, we previously discovered that *B. cinerea* delivers *Bc*sRNAs into plants to suppress host immunity genes [22, 43] most of them derive from LTR retrotransposons [21]. We here showed that natural *B. cinerea* strains that have lost production of LTR retrotransposon *Bc*sRNAs are less aggressive on tomato. We found that less aggressive *B. cinerea* strains failed to produce LTR retrotransposon *Bc*sRNAs possibly due to RIP. RIP on LTR retrotransposons is common in ascomycete species [30]; however, it remains unclear why some LTR retrotransposons escaped RIP in the aggressive *B. cinerea* isolate B05.10. This could be due to positive selection inferred by host plant infection. Moreover, transgene-induced production of retrotransposon *Bc*sRNAs in a non-retrotransposon *Bc*sRNAs producing *B. cinerea* strain led to enhanced aggressiveness in a *Bc*Gypsy3 RNA and *Bc*Gypsy3 BcsRNA expression-dependent manner. The level of transgene-induced *Bc*Gypsy3 *Bc*sRNA production in the recipient strain D08_H24 was dependent on the genome insertion site, but did not fully restore *Bc*sRNA production as found in the donor strain B05.10, although *BcGypsy3* mRNA expression levels were similar. Unexpectedly, there was a positive correlation between *Bc*Gypsy3 *Bc*sRNA expression and *Bc*Gypsy3 mRNA transcript levels. We speculate that post-transcriptional regulation might be the reason for moderate *Bc*sRNA production, or that stronger *Bc*sRNA production only occurs after several generations upon transformation [31, 44]. In addition, the retrotransposon transgene was transformed into D08_H24 without LTRs, which could be a hint that *Bc*sRNA production also depends on LTRs. Performing RNA-seq with *A. thaliana* infected with D08_H24 wt or a *Bc*Gypsy3-transformant, we revealed first insights into the regulatory effects of *B. cinerea* retrotransposon *Bc*sRNAs on the host plant transcriptome, comprising upregulation of immunity repressors as well as down-regulation of genes involved in activating immunity against *B. cinerea*. Since this analysis gave only a snapshot on plant gene manipulation induced by a single retrotransposon, we predict that more and other plant genes might be regulated through diverse LTR retrotransposon *Bc*sRNAs produced by the different *B. cinerea* isolates. Upregulation of LTR retrotransposon sRNAs was also evident in *Magnaporthe oryzae*, the fungal plant pathogen causing the rice blast disease, during infection of rice plants, as well as de-repression of LTR retrotransposon transcription was observed in the fungal wheat pathogen *Zymoseptoria tritici* [12] during wheat infection, suggesting a positive role of LTR retrotransposons in pathogenicity for other fungal pathogens. Moreover, animal-parasitic nematodes, such as *Heligmosomoides polygyrus*, secrete retrotransposon-derived sRNAs in extracellular vesicles [45], which are internalized by host cells and probably induce suppression of host mRNAs [46].

## Conclusion

In summary, we found evidence for correlation between pathogenicity and retrotransposons in a collection of natural fungal pathogen isolates. By a transgenic approach, we proved that retrotransposon expression in a weakly aggressive pathogen isolate significantly enhanced its aggressiveness level and that retrotransposons contribute to pathogenicity by producing trans-species sRNAs that manipulate host plant gene expression. It will be interesting to explore how common diverse pathogens and parasites utilize transposons as pathogenicity factors to colonize their hosts.

## Materials and Methods

### Strains materials, growth media and condition

The *Botrytis cinerea* (Pers.: Fr.) strain B05.10 as well as the five wild isolates D08_H24, D14_KF, D13_TF, D13_TS and N11_KW were used for this study. Routine cultivation was carried out on rich medium (RM; 10 g/L malt extract, 4 g/L yeast extract, 4 g/L glucose, 15 g/L agar) supplemented for mutant strains with hygromycin B (Sigma-Aldrich; 70 μg/ml) or nourseothricin (Werner Bioagents; 120 μg/ml). The plates were incubated on the bench in transparent plastic boxes under natural light conditions. *Arabidopsis thaliana* (L.) were grown on soil under short day condition (8 h light/ 16 h dark, 22 °C, 60 % relative humidity). *Solanum lycopersicum* (L.) (tomato) cultivars Moneymaker or Heinz were grown under 16 h light/8 h dark, 24°C, 60 % humidity condition.

### Pathogenicity assay

Pathogenicity assays were performed on detached tomato leaves from 4 to 5 weeks-old plants. *B. cinerea* conidia were resuspended in 1% malt extract at a final concentration of 5×10^4^ conidia/ml. Tomato leaves were inoculated with 20μl conidia solution or agar plugs from 4 days-old *Botryti*s mycelia grown on RM agar, and inoculated leaves were kept in a humidity box. Infected leaves were photographed and lesion area was measured using the Fiji software (ImageJ version 2.1.0/1.53c).

### Trypan Blue staining & microscopy

Infected tomato leaves were stained with Trypan Blue as described previously [47]. Microscopic images were taken with a DFC450 CCD-Camera (Leica) on a CTR 6000 microscope (Leica Microsystems). Microlesion quantification was done by the FIJI software by choosing the image-type 8-bit format (grey scale), scaling whole leaf diameter to 1, auto-setting of image adjust threshold and measure pixel counts in lesion area versus whole leaf area.

### Genotyping PCR

Genomic DNA was isolated using the CTAB method followed by chloroform extraction and isopropanol precipitation [48]. PCR primers used are given in Tab.S12.

### RNA extraction, Reverse transcription and quantitative PCR

Total RNA was isolated using a CTAB-based method [49]. Genomic DNA was removed using DNase I (Sigma-Aldrich) treatment and cDNA synthesis was performed with 1 μg total RNA using SuperScriptIII RT (Thermo Fisher Scientific). Gene expression was measured by qPCR on a Quantstudio5 cycler (ThermoFischer Scientific) using the Primaquant low ROX qPCR master mix (Steinbrenner Laborsysteme). Primers are listed in Tab.S12. *Bc*Tubulin was used for normalization. Differential expression was calculated using the 2^−ΔΔCt^ method [50].

### Stem-loop RT-PCR

Detection of *Bc*sRNAs was carried out following the stem-loop RT-PCR protocol [51] using 1 μg of total RNA. Primers are listed in Tab.S12. The stem-loop RT-PCR products were visualized on 10% non-denaturing polyacrylamide gels.

### Cloning and *B. cinerea* transformation

The different cloning constructs used for *B. cinerea* transformation were generated using the Golden Gate strategy [52] and are presented in Fig.S9A, and used primers are listed in Tab.S12. Before fungal transformation, the plasmids were digested with the restriction endonuclease BsaI (NEB Biolabs) to isolate the transformation cassette of interest. The transformation of *B. cinerea* strain D08_H24 was performed as described before [53] with the following minor modifications. Transformed protoplasts were mixed into SH agar without antibiotics and incubated in dark for 24 hours. Upon pre-incubation, SH agar with protoplasts were covered with fresh SH agar containing 70 μg/ml of hygromycin B (Sigma-Aldrich) or 120 μg/ml of nourseothricin (Werner Bioagents) and the plates were further incubated in the dark until the isolation of putative mutants. Transformants with targeted insertion of transgenes into the *BcniaD* locus were further selected by plating on Czapek-Dox agar containing 0.4 M potassium chlorate as previously described [33]. Successful target insertion was confirmed by genomic PCR analysis.

### Illumina sRNA-seq and data analysis

sRNAs were isolated from *B. cinerea* grown on RM for 4 days for high throughput sequencing, as previously described [22]. sRNAs were cloned for Illumina sequencing using the Next® Small RNA Prep kit (NEB) and sequenced on an Illumina HiSeq1500 platform. The Illumina sequencing data were analysed using the GALAXY Biostar server [54]. Raw data were de-multiplexed (Illumina Demultiplex, Galaxy Version 1.0.0) [40] and adapter sequences were removed (Clip adaptor sequence, Galaxy Version 1.0.0). Reads were mapped to *B. cinerea* B05.10 reference genome assembly (BioProject: PRJNA15632) or *de-novo* genome assemblies of the strains D08_H24 and D14_KF using the BOWTIE algorithm (Galaxy Version 1.1.0) allowing zero mismatches (-v 0). Ribosomal RNAs (rRNA) were filtered out using the BOWTIE algorithm allowing three mismatches (-v 3). Reads were sorted to transfer RNA (tRNA), small nuclear/nucleolar RNA (snoRNA), messenger RNA, and transposon RNA reads using BOWTIE2 [55] with default settings. Upon classification, reads were count and normalized on total *B. cinerea* reads per million (RPM). Mapping files were visualized using a custom script based on ggplot2 (version 3.2.1) package in R (version 3.6.1). Target gene prediction of sRNAs was performed with the TAPIR program using a maximal score of 4.5 and a free energy ratio of 0.7 as thresholds [56] and psRNATarget with default settings [40].

### RNA-seq

Conidia from 10-day-old mycelium cultures were harvested and diluted to 10^6^ conidia/ml in 1% malt extract (pH 5.7). Six-week-old *Arabidopsis thaliana* Col-0 plants were infected with those conidia (24 plants / strain dispatched into 2 humidity boxes). For each plant, 5 leaves were infected with 4 drops of 5 μl. After 24 hours incubation on the bench, 3 leaves from each plant were randomly harvested. Leaves containing the 4 spots of infection were cut and the infections from 2 leaves were pooled into a single Eppendorf tube and snap frozen in liquid nitrogen constituting one biological replicate. Total RNAs were extracted using a CTAB-based method [49]. Total RNA was subjected to mRNA sequencing using a version of the prime-seq method online available at https://www.protocols.io/view/prime-seq-s9veh66. This protocol is based on single cell RNA-seq [57] and is a three-prime counting method that includes a sample specific barcode sequence and unique molecular identifiers (UMI) for accurate quantification of gene expression. Illumina paired end sequencing was performed on an HiSeq 1500 instrument. Raw data was demultiplexed using deML [58], adapters and poly-A tails were trimmed using cutadapt (version 2.3) and further preprocessed using the zUMIs pipeline [59] with STAR [60]. Reads were mapped to combined *A. thaliana* (TAIR10) or *B. cinerea* (ASM83294v1) genomes with Araport11 and ASM83294v1.41 gene annotations. Differential gene expression analysis was done in iDEP.91 [61] with FDR cut-off at 0.05 using limma-voom normalization.

### Whole genome sequencing

WGS of *B. cinerea* D08_H24 and D14_KF strains was accomplished by hybrid Oxford Nanopore (Promethion) long-read sequencing and Illumina HiSeq1500 short-read sequencing. After base calling with guppy (version 2.3.7) and adapter removal with porechop (https://github.com/rrwick/Porechop), the long-reads where assembled using wtdbg2 [62] and polished with two rounds of racon [63]. The short reads were used to polish the assembly with pilon [64]. The Circos plotting library [65] was used via R circlize [66] to visualise whole genome comparisons.

### Transposon annotation and RIP analysis

Transposable elements in the *B. cinerea* strain B05.10 were previously annotated [23] and used in this study. Annotation of transposable elements in D08_H24 and D14_KF genomes was performed using the REPET package as described previously [23]. Sequence alignment and phylogenetic tree generation was done using the CLC main workbench software (version 20.0.4, Qiagen) using the Neighbor Joining method and Jukes-Cantor nucleotide distance measure. Bootstrap analysis was run with 500 replicates. Blastn search was used to identify truncated retrotransposons (> 400 bp) with the following optional parameters -word_size 20 - max_target_seqs 50000 -gapopen 5 -gapextend 2 -reward 2 -penalty −3 -dust no - soft_masking false. The GC richest copy of each LTR retrotransposon subfamilies in B05.10 was chosen as a query. The sequences of the hits longer than 1000 bp were extracted and aligned with the refalign tool from the REPET package (https://urgi.versailles.inra.fr/Tools/REPET) to further run RIPCAL [29] analysis using the GC-richest query copy as a reference. RIPCAL was used to calculate all possible RIP mutations (CA → TA, CT → TT, CC → TC, CG → TG and their reverse complements). For the calculation of the RIP index TA/AT [67], only full-length copies from the REPET analysis were used. The different dinucleotide combinations were counted with the compseq tool (from the EMBOSS package) and used to calculate the RIP index.

Prediction of protein domains in LTR retrotransposons was done on a GC-rich copy sequence of each consensus class using the NCBI conserved domain database [68]. Long terminal repeats sequence and size were obtained from previous analysis [23]. For *Bc*Gypsy1, the additional ORF sequence (*brtn*) was obtained from Zhao et al. [69]. It is to note that this gene is not found in the last annotation of B05.10 genome. For the P26.1 consensus, the beginning of *brtn* is not present as in the other consensus grouped as the *Bc*Gypsy1 subfamily but a putative ORF is still present (dashed arrow in Fig.S1). The other represented additional ORFs were predicted in the last annotation of B05.10 genome (http://fungi.ensembl.org/Botrytis_cinerea). The predicted protein domain figure was generated using IBS (http://ibs.biocuckoo.org/).

### Plots & statistical analysis

Plots and statistical tests were carried out using R studio (version 1.0.136, rstudio.com), ggplot2 (version 3.2.1) or Excel.

## Acknowledgements

The authors thank Verena Klingl for excellent technical assistance and Dr. Andreas Brachmann as well as the Gene Center Munich for Nanopore and Illumina sequencing service. We further would like to thank Dr. Claude Becker and Florian Dunker for manuscript proofreading and helpful suggestions. We also thank Dr. Martin Parniske and Dr. David Chiasson for providing us Golden Gate entry plasmids. We thank Adeline Simon for sharing the compseq script and for bioinformatics assistance and the members of URGI /INRAE in Versailles for the manual curation of transposable elements annotations workshop and for server access. A.P. was supported by the Alexander von Humboldt Foundation (individual postdoctoral research fellowship) and the project was supported by Sonderforschungsbereich SFB924 (Grant ID: INST 95/1552-1). The funders had no role in study design, data collection and analysis, and decision to publish or in preparation of the manuscript.

## Author contributions

Research concept and design A.W.; experimental design: A.P., and A.W.; experiments performed A.P, C.T., C.G.; RNA-seq A.P. L.W. W.E.; WGS A.P., A.H.; bioinformatics analysis: A.W., A.P., F.S.; manuscript writing: A.W. and A.P.; manuscript reviewing and editing: A.P., C.T., F.S., M.H. and A.W.

## Availability of data and materials

All small RNA-seq, WGS, and RNA-seq raw data will be submitted to NCBI SRA repository.

## Supplemental material

**Figure S1:** Conserved domains of LTR retrotransposons in B05.10 representing the nine consensus classes and the six subfamilies.

**Figure S2:** A) Copy number, nucleotide count, and mapped *Bc*sRNAs of the six LTR retrotransposon subfamilies *Bc*Gypsy1-4, *Bc*Copia1-2. B) *Bc*sRNA maps at the *BcGypsy1*/*BcGypsy3* locus at different time points of *A. thaliana* or *S. lycopersicum* host infection. Bars above line represents sense and below line antisense reads. C) *Bc*sRNA abundance in reads per million total *Bc*sRNAs mapping to tRNA, sn/snoRNA or mRNA gene loci at different time points of *S. lycopersicum* or *A. thaliana* host infection, with 0 hpi implies *B. cinerea* spore inoculation and direct sample harvest. For B) and C), raw data are available at NCBI GEO: GSE45323, GSE45321.

**Figure S3:** A) Radial growth speed and B) sclerotia formation of the six *B. cinerea* strains. Scale bar in B) represents 2 cm. For A), Error bars represent standard deviation from three biological replicates.

**Figure S4:** *Bc*sRNAs of the six *B. cinerea* strains mapping to a *Bc*Gypsy1/*Bc*Gypsy3 locus in B05.10 exemplifying reads aligned in sense and antisense are represented in blue and red, respectively, and the bar plots represent the percentage of reads with a specific nt length. Different RPM scales are shown; 275 RPM for N11_KW, 500 RPM for B05.10, 20 RPM D13_TF, 1 RPM for D14_KF and 2 RPM for D13_TS and D08_H24.

**Figure S5:** A) Genome coverage of LTR retrotransposon subfamilies in the strains B05.10, D08_H24 and D14_KF. B) Copy length distribution of LTR retrotransposons in the strains B05.10, D08_H24 and D14_KF.

**Figure S6:** Analysis of GC content (%) of truncated LTR retrotransposons identified by Blastn search.

**Figure S7:** RIPCAL analysis of the of the *Bc*Gypsy3 subfamily in the strains B05.10, D08_H24 and D14_KF. Similar results were found for all six LTR retrotransposon families, *Bc*Gypsy1-4, *Bc*Copia1, *Bc*Copia2. Each line represents one *Bc*Gypsy3 copy. The GC-richest *Bc*Gypsy3 copy of B05.10 was used as reference for alignment with *Bc*Gypsy3 copies in D08_H24 and D14_KF.

**Figure S8:** BLASTn search of *BcGypsy3* in B05.10 and D08.H24

**Figure S9:** A) Cloning strategy and B) genotyping PCR of the integrated *BcGypsy3* transgene into the *BcniaD* locus. M: 1 kb DNA ladder marker. C) Morphological phenotypes of D08_H24 *Bc*Gypsy3 transformants.

**Figure S10:** *Bc*sRNA size profiles and 5`end first nucleotide distribution of *Bc*Gypsy3 transformants.

**Figure S11:** A-B) Sequence alignment of *Bc*sRNA effectors to *S. lycopersium* (*Sl*) or *A. thaliana* (*At*) mRNA target candidates. C) qRT-PCR analysis of *A. thaliana* target mRNAs after no-infection (mock) or after infection with D08_H24 wt or rand*Bc*Gypsy3 #1. *AtPDF1.2* and *AtPR1* were used as *B. cinerea*-inducible genes in *A. thaliana* and these gene were not predicted targets of *Bc*sRNAs. Each data point represents a biological replicate.

**Figure S12:** Change-fold factors of expressed genes found in *A. thaliana* mock versus *B. cinerea* infection, according to the BAR database.

**Table S1:** LTR retrotransposon consensus classes and subfamilies in *B. cinerea* B05.10.

**Table S2:** Class I and class II transposons of the strain B05.10.

**Table S3:** Host and geographical origin of the six *B. cinerea* strains.

**Table S4:** Raw reads of *Bc*sRNAs detected by sRNA-seq analysis of the six *B. cinerea* strains.

**Table S5:** Full-length LTR retrotransposons in the strains D08_H24 and D14_KF.

**Table S6:** Mapping analysis of *Bc*sRNAs in the strains B05.10, D08_H24 and D14_KF using the re-sequenced genome and annotated full-length and truncated LTR retrotransposons identified by REPET and Blastn search.

**Table S7:** Dinucleotide ratios of full-length LTR retrotransposons identified by REPET in the strains B05.10, D08_H24 and D14_KF.

**Table S8:** Correlation analysis of LTR retrotransposon *Bc*sRNA expression to GC content (%) and TA/AT ratio in B05.10.

**Table S9:** *Bc*sRNA mapping values at full-length LTR retrotransposons in B05.10.

**Table S10:** Relative expression values from RNA-seq analysis of differentially expressed *A. thaliana* genes infected with D08_H24 wt or rand*Bc*Gypsy3 #1.

**Table S11:** Predicted target genes of *Bc*Gypsy3 *Bc*sRNAs.

**Table S12:** DNA oligonucleotides used in this study.

