## Supplemental Figures for "Retrotransposons as pathogenicity factors of the plant pathogenic fungus *Botrytis cinerea*"

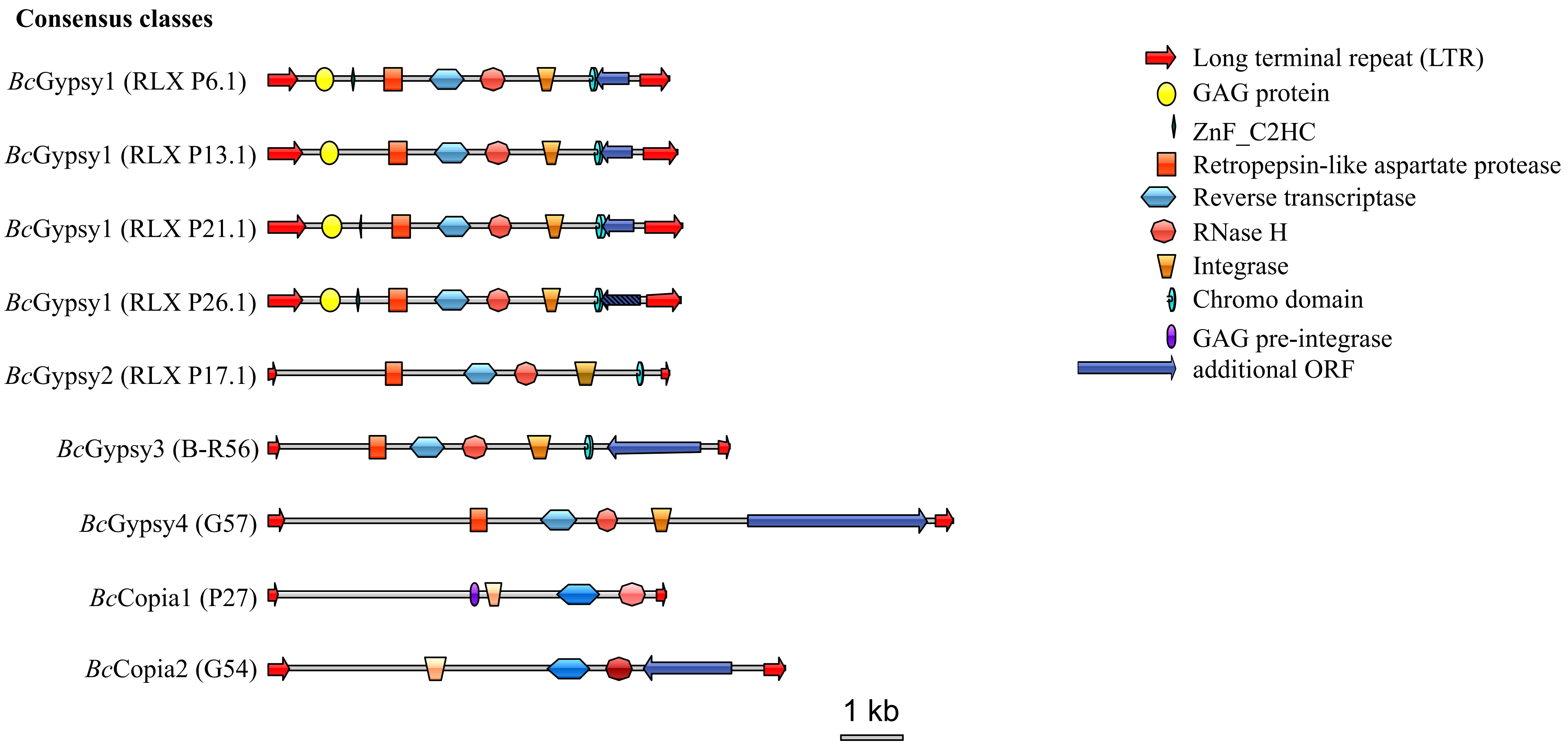

Consensus classes
BcGypsy1 (RLX P6.1)
BcGypsy1 (RLX P13.1)
BcGypsy1 (RLX P21.1)
BcGypsy1 (RLX P26.1)
BcGypsy2 (RLX P17.1)
BcGypsy3 (B-R56)
BcGypsy4 (G57)
BcCopia1 (P27)
BcCopia2 (G54)
1 kb
Long terminal repeat (LTR)
GAG protein
ZnF_C2HC
Retropepsin-like aspartate protease
Reverse transcriptase
RNase H
Integrase
Chromo domain
GAG pre-integrase
additional ORF


Figure S1

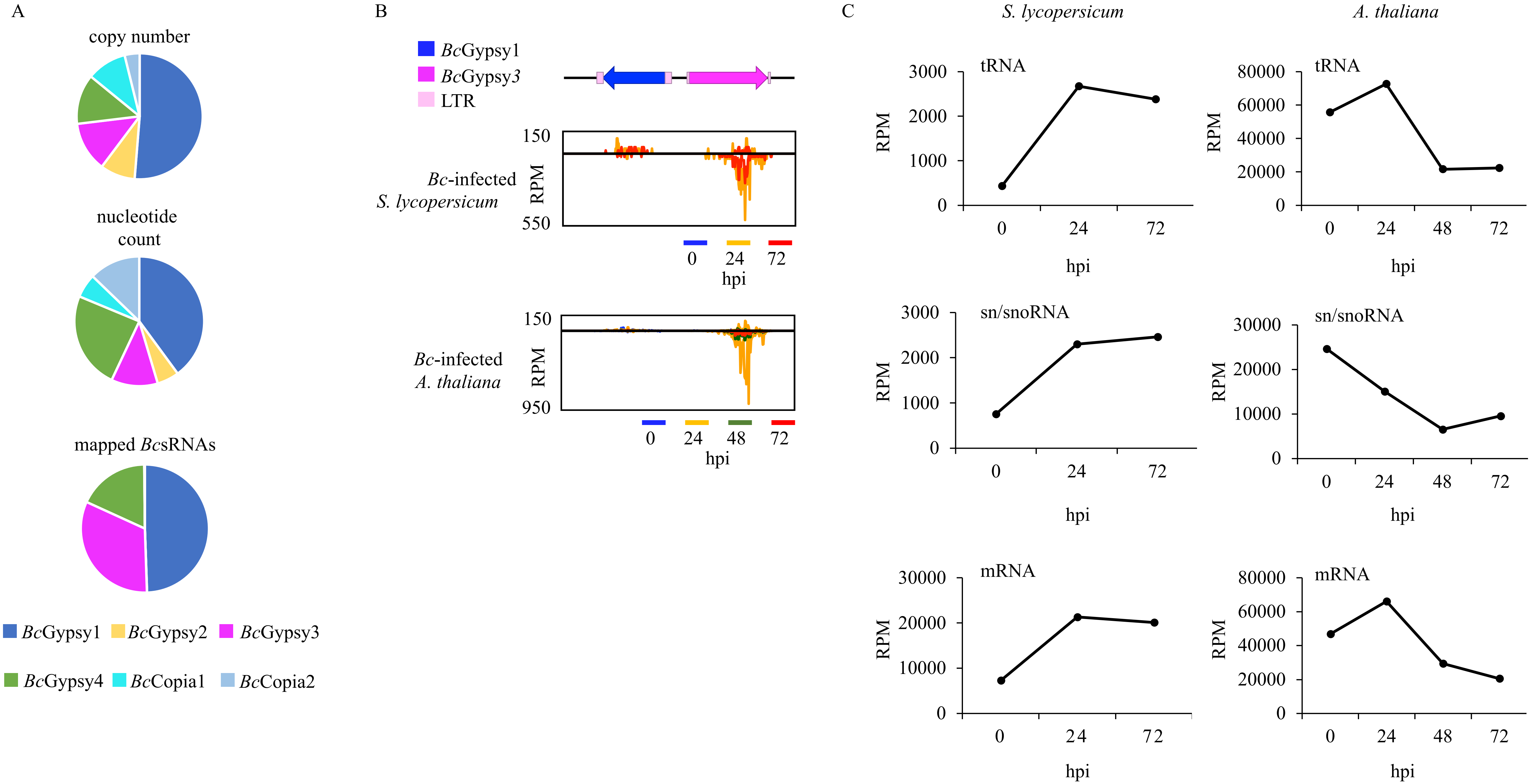

A
copy number
nucleotide count
mapped BcsRNAs
BcGypsy1
BcGypsy2
BcGypsy3
BcGypsy4
BcCopia1
BcCopia2
B
BcGypsy1
BcGypsy3
LTR
Bc-infected S. lycopersicum
RPM
150
550
0
24
72
hpi
Bc-infected A. thaliana
RPM
150
950
0
24
48
72
hpi
C
S. lycopersicum
A. thaliana
tRNA
sn/snoRNA
mRNA
RPM
hpi
0
24
48
72
0
1000
2000
3000
10000
20000
30000
40000
60000
80000


Figure S2

A

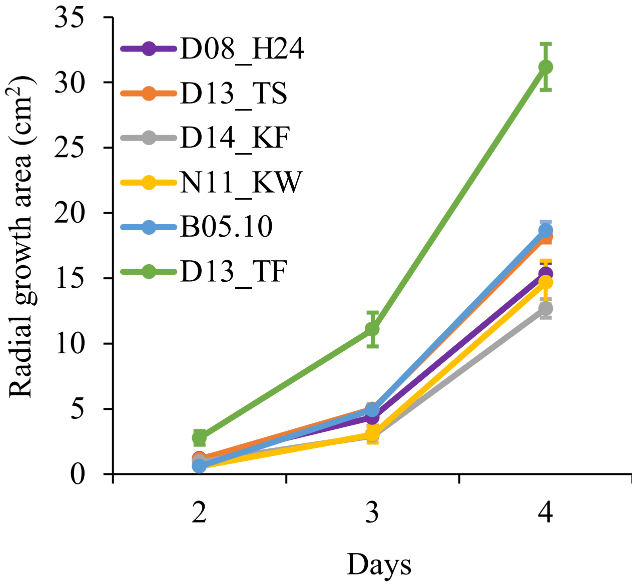

35
30
25
20
15
10
5
0
Radial growth area (cm$^2$)
2
3
4
Days
D08_H24
D13_TS
D14_KF
N11_KW
B05.10
D13_TF


B

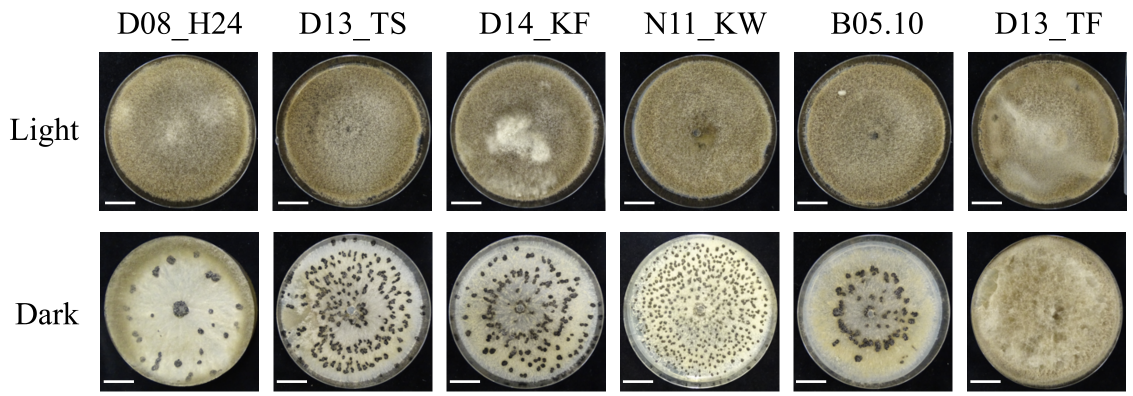

D08_H24
D13_TS
D14_KF
N11_KW
B05.10
D13_TF
Light
Dark


Figure S3

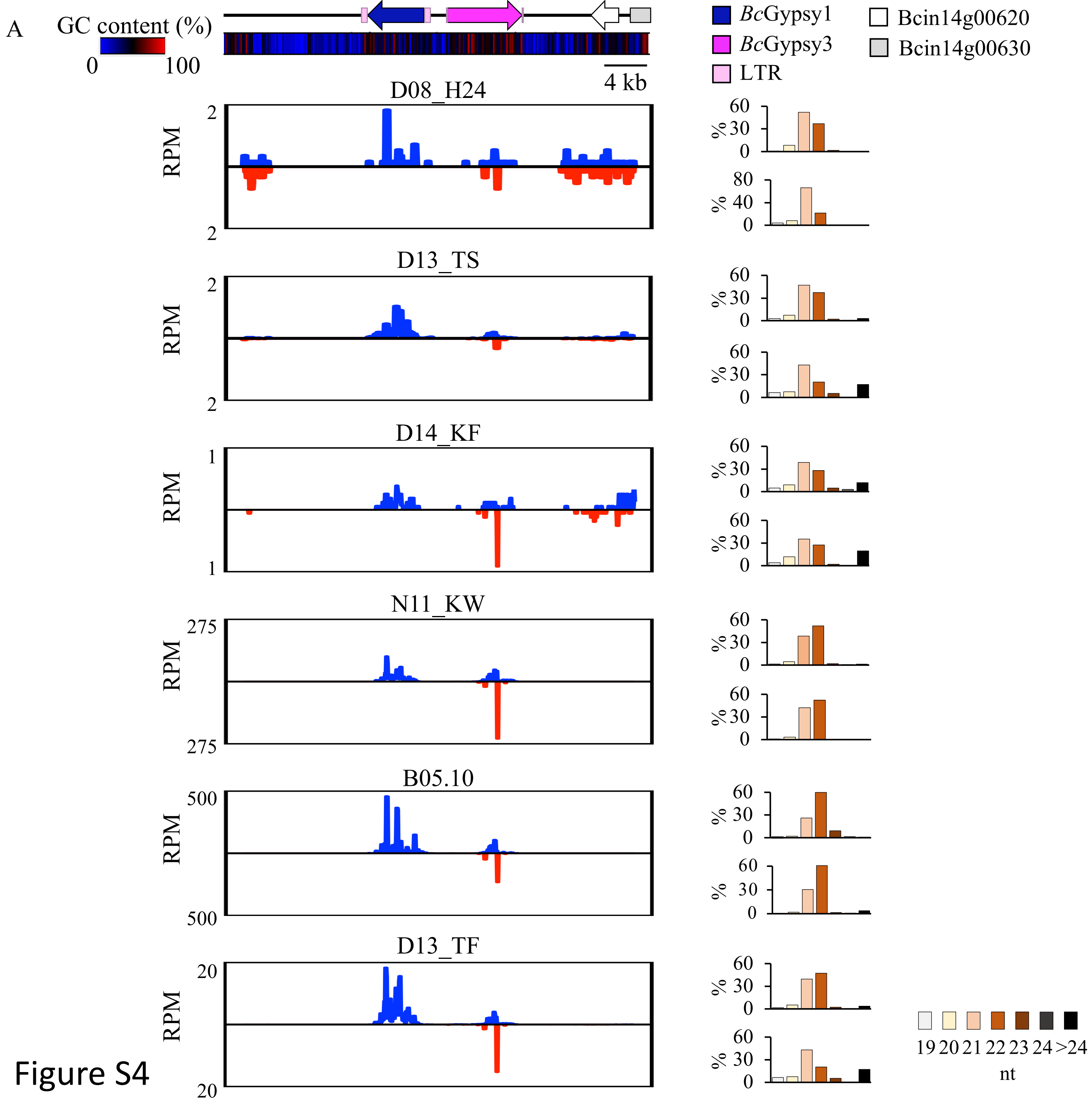
A
GC content (%)
0
100
4 kb
BcGypsy1
BcGypsy3
LTR
Bcin14g00620
Bcin14g00630
D08_H24
D13_TS
D14_KF
N11_KW
B05.10
D13_TF
RPM
%
19 20 21 22 23 24 >24
nt

Figure S4

A

| | B05.10 | D08_H24 | D14_KF |
|---|---|---|---|
| *Bc*Gypsy1 | 357,565 bp<br>0.84 % | 92,081 bp<br>0.22 % | 164,229 bp<br>0.38 % |
| *Bc*Gypsy2 | 49,141 bp<br>0.12 % | 21,556 bp<br>0.05 % | 64,674 bp<br>0.15 % |
| *Bc*Gypsy3 | 103,916 bp<br>0.24 % | 7,952 bp<br>0.02 % | 37,056 bp<br>0.09 % |
| *Bc*Gypsy4 | 217,377 bp<br>0.51 % | 45,201 bp<br>0.11 % | 77,969 bp<br>0.18 % |
| *Bc*Copia1 | 53,275 bp<br>0.13 % | 31,145 bp<br>0.07 % | 7,152 bp<br>0.02 % |
| *Bc*Copia2 | 114,550 bp<br>0.27 % | 22,194 bp<br>0.05 % | 70,416 bp<br>0.16 % |
| **Total coverage** | **895,824 bp**<br>**2.10 %** | **220,129 bp**<br>**0.52 %** | **421,496 bp**<br>**0.99 %** |

B

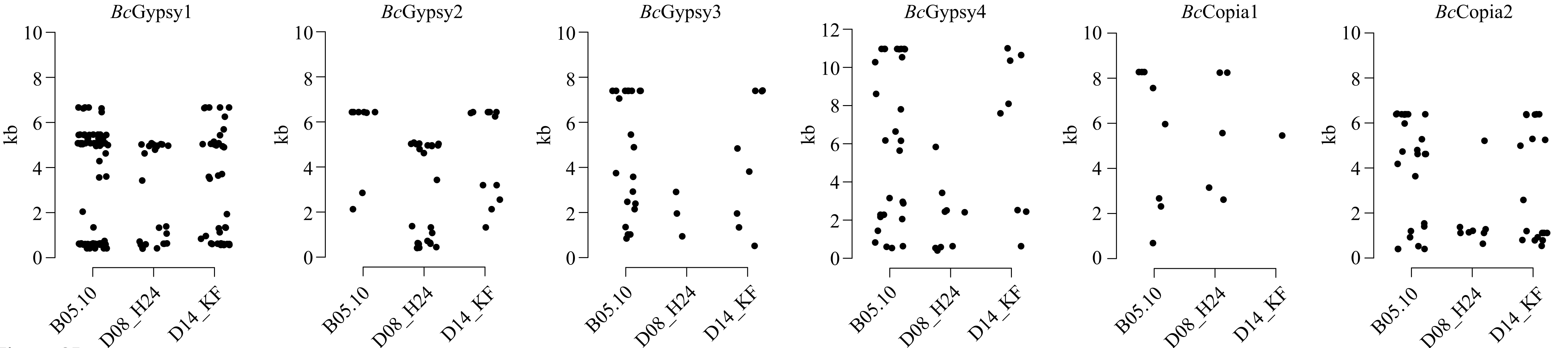


Figure S5

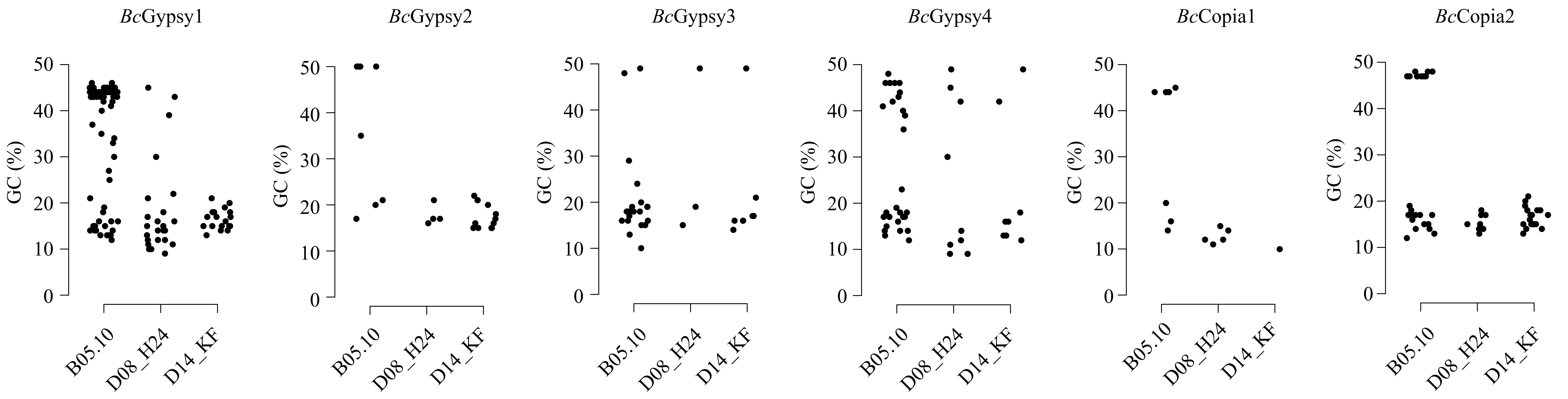

BcGypsy1
BcGypsy2
BcGypsy3
BcGypsy4
BcCopia1
BcCopia2
GC (%)
0
10
20
30
40
50
B05.10
D08_H24
D14_KF


Figure S6

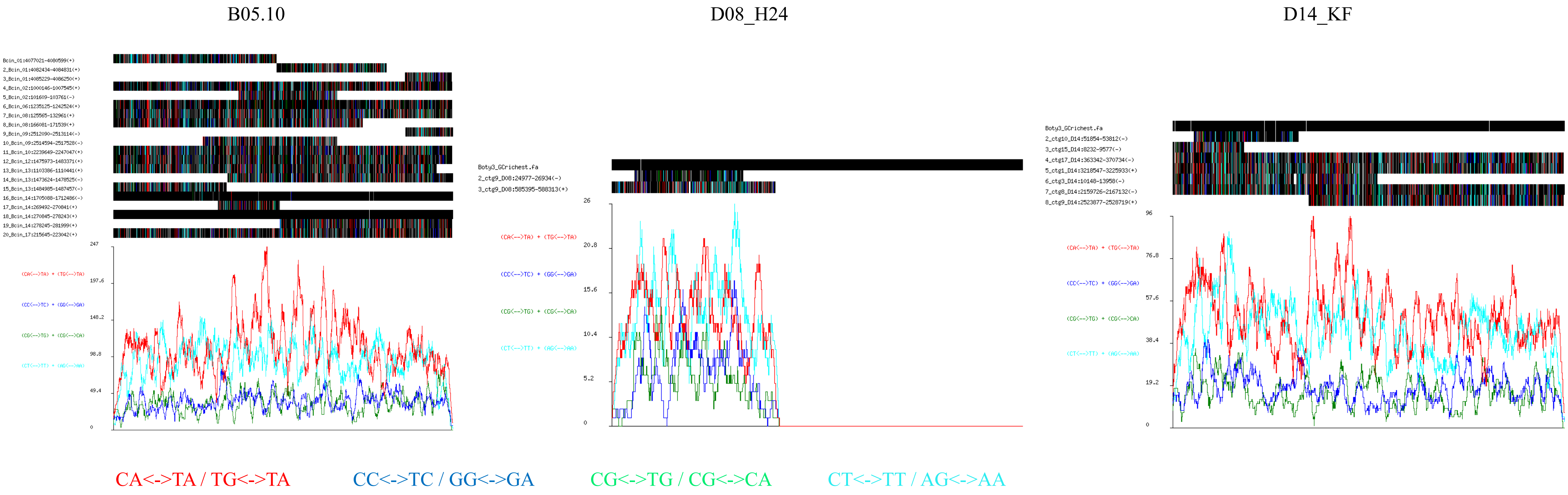


Figure S7

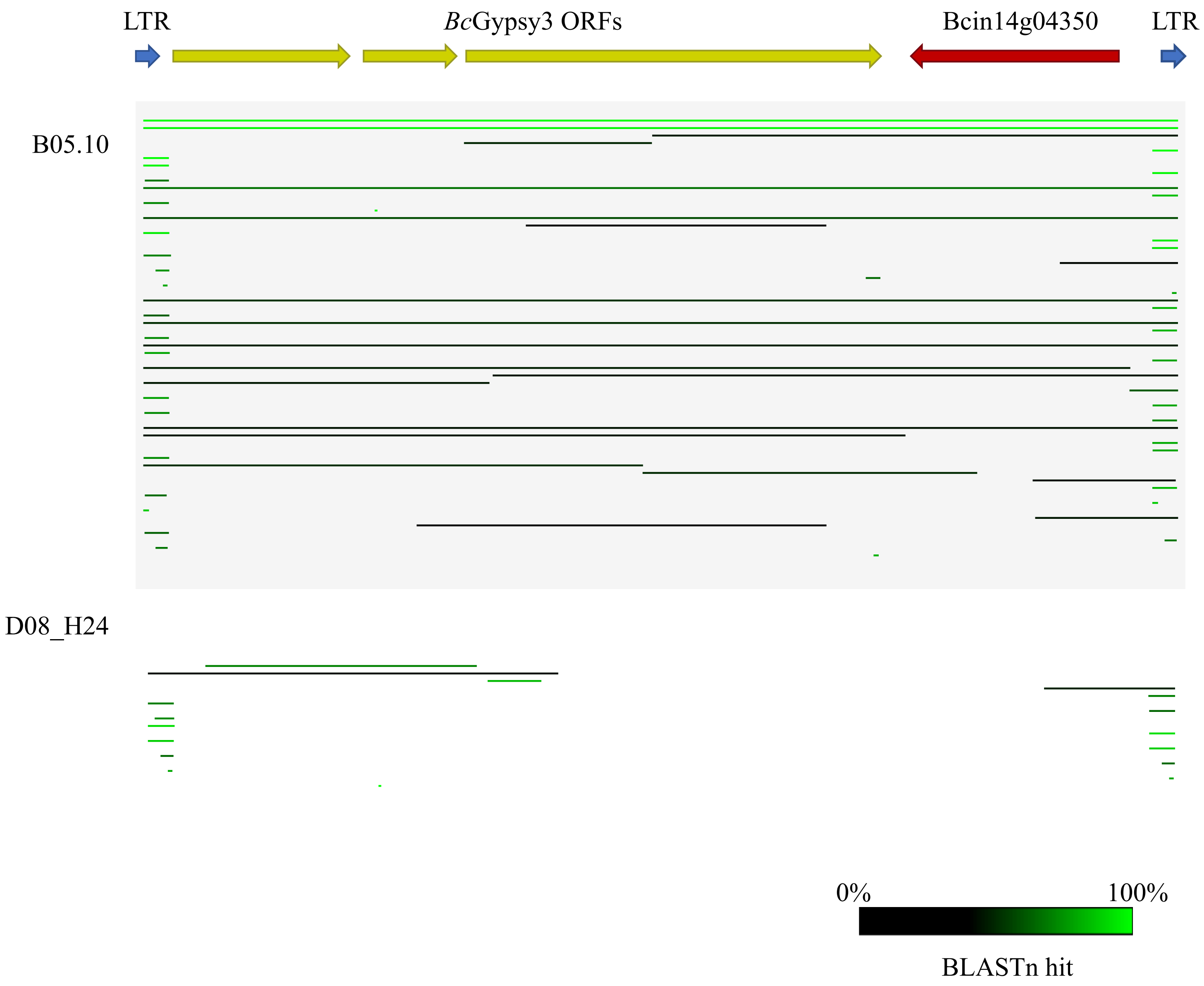
LTR
BcGypsy3 ORFs
Bcin14g04350
LTR
B05.10
D08_H24
0%
100%
BLASTn hit

Figure S8

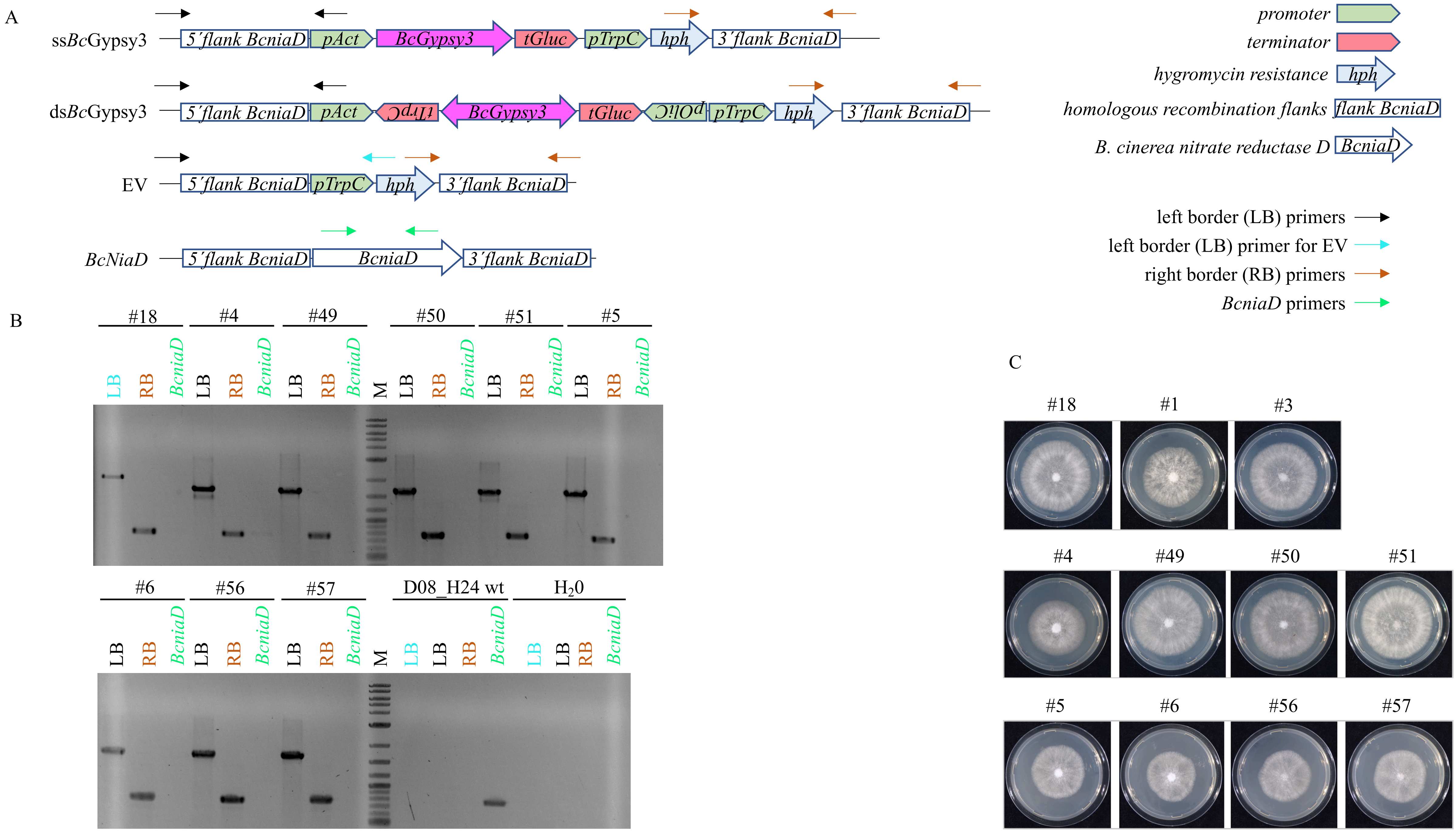
A
ssBcGypsy3
5´flank BcniaD
pAct
BcGypsy3
tGluc
pTrpC
hph
3´flank BcniaD
dsBcGypsy3
5´flank BcniaD
pAct
tTrpC
BcGypsy3
tGluc
pOliC
pTrpC
hph
3´flank BcniaD
EV
5´flank BcniaD
pTrpC
hph
3´flank BcniaD
BcNiaD
5´flank BcniaD
BcniaD
3´flank BcniaD
promoter
terminator
hygromycin resistance
hph
homologous recombination flanks
flank BcniaD
B. cinerea nitrate reductase D
BcniaD
left border (LB) primers
left border (LB) primer for EV
right border (RB) primers
BcniaD primers
B
#18
#4
#49
#50
#51
#5
LB RB BcniaD LB RB BcniaD LB RB BcniaD M LB RB BcniaD LB RB BcniaD LB RB BcniaD
#6
#56
#57
D08_H24 wt
H20
LB RB BcniaD LB RB BcniaD LB RB BcniaD M LB LB RB BcniaD LB LB RB BcniaD
C
#18
#1
#3
#4
#49
#50
#51
#5
#6
#56
#57

Figure S9

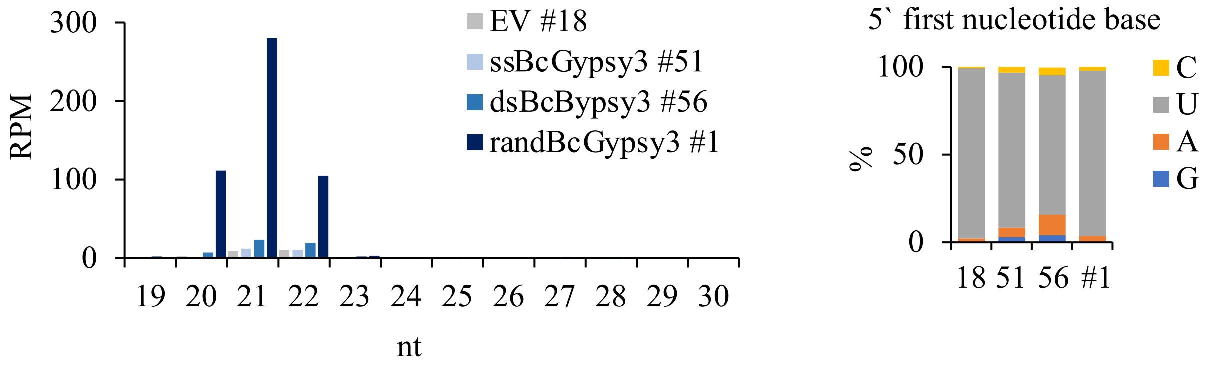
300
200
100
0
RPM
19 20 21 22 23 24 25 26 27 28 29 30
nt
EV #18
ssBcGypsy3 #51
dsBcBypsy3 #56
randBcGypsy3 #1
5` first nucleotide base
100
50
0
%
18 51 56 #1
C
U
A
G

Figure S10

A

```
BcsRNA3.2 3'GUGGAUGUUCUAGGUGUUACA 5'
            ||::|| ||||||||| ||||
SlMPKKK4  5'CAUUUAAAAGAUCCACCAUGU 3'

BcsRNA3.1 3'CGGGUGGAUGUUCUAGGUGUU 5'
              |||||||:||| ||||||:|
Slhse1    5'ACCCACCUGCAACAUCCACGA 3'

BcsRNA20 3'UUAGUCUUUUUGUUCUUGUGAU 5'
           |:| |||||||||| ||:||||
SlBhlh63 5'AGUAAGAAAAACAAUAAUACUA 3'
```

B

```
BcsRNA3.2 3'UGGAUGUUCUAGGUGUUACAU 5'
            |:| | |||||:||||||||
AtMPK1    5'AUCAAGAAGAUUCACAAUGUU 3'

BcsRNA3.2 3'UGGAUGUUCUAGGUGUUACAU 5'
            |:| | ||||||||||||||:
AtMPK2    5'AUCAAGAAGAUCCACAAUGUG 3'

BcsRNA3.1 3'CGGGUGGAUGUUCUAGGUGUU 5'
            |||:| |||||||| |||||
AtPRXIIF  5'GCCUAGCUACAAGAGCCACAU 3'
```

C

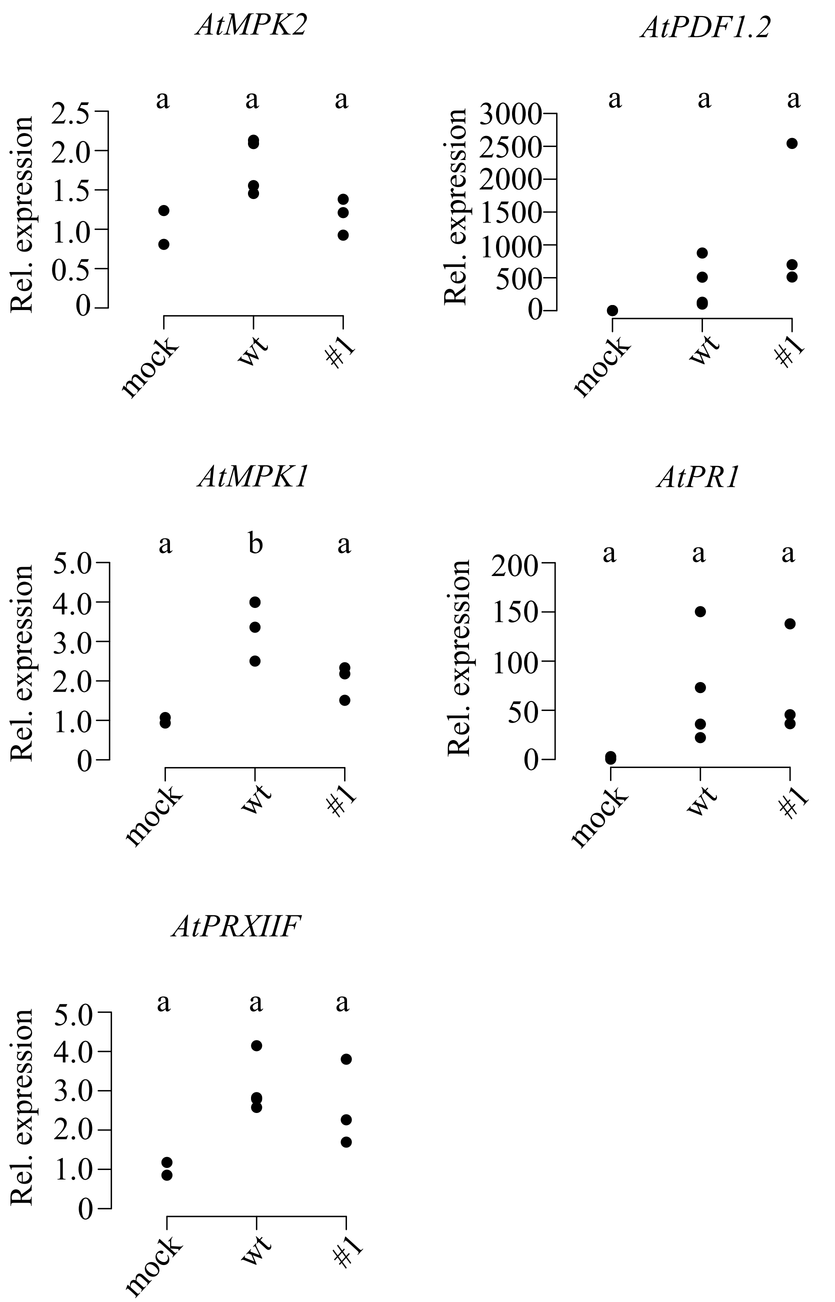


Figure S11

| | mock | | *B. cinerea* B05.10 | | |
|---|---|---|---|---|---|
| | 18 | 48 | 18 | 48 | hpi |
| *AT1G56090* | 1 | 1 | 0.7 | 0.8 | |
| *AT3G01150* | 1 | 1 | 0.9 | 0.9 | |
| *AT1G08465* | n.d. | n.d. | n.d. | n.d. | |
| *AT4G12830* | 1 | 1 | 0.7 | 0.5 | |
| *AT1G72430* | 1 | 1 | 0.6 | 0.5 | |
| *AT1G52870* | 1 | 1 | 0.7 | 0.6 | |
| *AT1G47128* | 1 | 1 | 0.8 | 0.3 | |
| *AT4G35090* | 1 | 1 | 0.7 | 0.2 | |
| *AT1G77060* | 1 | 1 | 0.7 | 0.5 | |
| *AT1G10170* | 1 | 1 | 0.7 | 0.7 | |
| *AT1G08050* | 1 | 1 | 3.5 | 10.9 | |
| *AT1G17380* | 1 | 1 | 1.1 | 1.9 | |
| *AT2G43150* | 1 | 1 | 1 | 1.5 | |
| *AT3G10985* | 1 | 1 | 1.3 | 1.4 | |
| *AT4G24350* | 1 | 1 | 4.8 | 3.4 | |

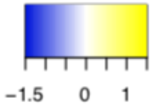


Figure S12
